# Regulatory variants explain much more heritability than coding variants across 11 common diseases

**DOI:** 10.1101/004309

**Authors:** Alexander Gusev, S. Hong Lee, Benjamin M. Neale, Gosia Trynka, Bjarni J. Vilhjálmsson, Hilary Finucane, Han Xu, Chongzhi Zang, Stephan Ripke, Eli Stahl, Schizophrenia Working Group of the Psychiatric Genomics Consortium, SWE-SCZ Consortium, Anna K. Kähler, Christina M. Hultman, Shaun M. Purcell, Steven A. McCarroll, Mark Daly, Bogdan Pasaniuc, Patrick F. Sullivan, Naomi R. Wray, Soumya Raychaudhuri, Alkes L. Price

## Abstract

Common variants implicated by genome-wide association studies (GWAS) of complex diseases are known to be enriched for coding and regulatory variants. We applied methods to partition the heritability explained by genotyped SNPs 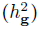 across functional categories (while accounting for shared variance due to linkage disequilibrium) to genotype and imputed data for 11 common diseases. DNaseI Hypersensitivity Sites (DHS) from 218 cell-types, spanning 16% of the genome, explained an average of 79% of 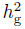 (5.1× enrichment; *P* < 10^−20^); further enrichment was observed at enhancer and cell-type specific DHS elements. The enrichments were much smaller in analyses that did not use imputed data or were restricted to GWAS-associated SNPs. In contrast, coding variants, spanning 1% of the genome, explained only 8% of 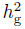 enrichment; *P* = 5 × 10^−4^). We replicated these findings but found no significant contribution from rare coding variants in an independent schizophrenia cohort genotyped on GWAS and exome chips.

---

Recent work by ENCODE and other projects has shown that specific classes of non-coding variants can have complex and diverse impacts on cell function and phenotype^1–7^. With many potentially informative functional categories and competing biological hypotheses, quantifying the contribution of variants in these categories to complex traits would inform trait biology and focus fine-mapping. The availability of significantly associated variants from hundreds of genome-wide association studies (GWAS)^8^ has opened one avenue for quantifying enrichment. Indeed, 11% of GWAS hits lie in coding regions^8^ and 57% of GWAS hits lie in broadly-defined DHS (spanning 42% of the genome)^5^, with additional GWAS hits tagging these regions. The full distribution of GWAS association statistics exhibits enriched P-values in coding and untranslated regions (UTR)^9^. Analysis of DHS sub-classes and other histone marks has revealed a complex pattern of cell-type specific relationships with known disease associations^4^. However, the question of how much each functional category contributes to disease heritability remains unanswered^10^.

Here, we jointly estimate the heritability explained by all SNPs 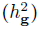 in different functional categories, generalizing recent work using variance-component methods^11–17^. In contrast to analyses of top GWAS hits, this approach leverages the entire polygenic architecture of each trait and can obtain accurate estimates even in the face of pervasive linkage disequilibrium (LD) across functional categories, as we show via extensive simulations. We apply this approach to functional categories in GWAS and exome chip data from > 50, 000 samples.

## Results

### Overview of methods

We applied the variance-components approach for estimating 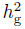^11,14^ to multiple components, each constructed from variants belonging to a functional category. The underlying model assumes that SNP effect-sizes are drawn from a normal distribution with category-specific variance, and relates the observed phenotypic covariance to a weighted sum of genetic relationship matrices computed from SNPs in each category. The joint estimate allows all components to compete for shared variance between categories due to LD. We confirmed by extensive simulations using real genotypes and functional categories that this approach achieves robust estimates under diverse disease architectures (see Methods).

We annotated the genome using six primary categories (see Methods, Table S1): (1) coding, (2) untranslated region (UTR), (3) promoter, (4) DHS in any of 218 cell-types, (5) intronic, and (6) intergenic. We then assigned each SNP a unique annotation defined by the first of these categories that it was annotated with, resulting in six non-overlapping variance-components (with the DHS category thus restricted to distal regions). Each resulting category exhibits similar average allele frequency and imputation accuracy (Table S2). After jointly estimating the variance explained by each category, we rescaled these values (and their standard errors) to the fraction of total 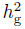 explained and compared this with category size (fraction of SNPs) to quantify enrichment (see Methods).

### Heritability of functional categories across 11 diseases

We analyzed a total of 11 WTCCC and WTCCC2 phenotypes^18–20^. After quality control^16^, the 7 WTCCC traits each included an average of 1,700 cases and a set of 2,700 shared controls; the 4 WTCCC2 traits included 1,800-9,300 cases and 5,300 shared controls (see Methods, Table S3). In all analyses of autoimmune traits, SNPs in the well-studied MHC region were excluded, though inclusion of the MHC as a separate component did not significantly affect the results. Each cohort was imputed to the 1000 Genomes reference panel yielding 4-6 million SNPs per trait after QC (see Methods, Table S3).

Combined results meta-analyzed across all traits are displayed in Figure 1 (Table S4). When including imputed SNPs, variants in DHS exhibited the greatest 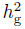 and most significant enrichment, with imputed DHS SNPs explaining an average of 79% (s.e. 7%) of the total 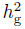, a 5.1 × enrichment (*P* < 10^−20^); the enrichment varied across traits (Fig. S1, Table S5), with a nominally significant difference between the 6 autoimmune (AS, CD, MS, RA, T1D, UC) and 5 non-autoimmune traits (5.5× and 3.3×; *P* = 0.01). Coding variants exhibited the greatest overall enrichment at 13.8× (*P* = 4.8 × 10^−4^) but accounted for a small fraction of 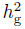 due to the much smaller category size. Correspondingly, we observed a significant depletion for both intronic regions (0.1×; *P* = 5.7 ×; 10^−12^) and intergenic regions (−0.1×; *P* < 10^−20^), with 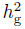 not significantly different from zero. Although individual values of 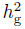 were allowed to fluctuate outside the 0-1 bound on variance to achieve unbiased estimates, a constrained analysis yielded similar results (see Table S7). After narrowing the DHS peaks to cover only 1% of the genome, the resulting DHS component still directly explained 20% (s.e. 4%, *P* = 2.9 × 10^−6^) of the total 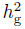, and explained 61% in a univariate model (which includes tagging of all functional categories; see Supplementary Note, Table S6). All enrichments/depletions were much weaker and less informative (measured by information content, see Supplementary Note) when restricting to genotyped SNPs, consistent with our simulations (see Methods, Fig. 1).

**Figure 1.**
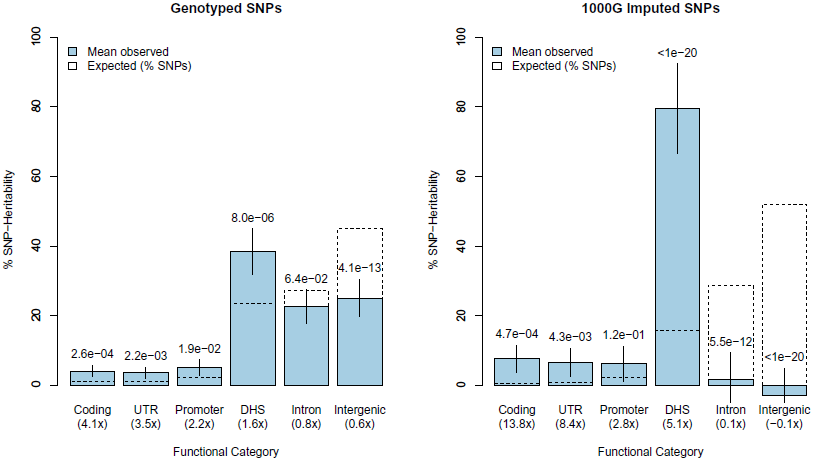
Functional partitioning of SNP-heritability across eleven traits. For each trait, the liability explained by SNPs in six functional categories was jointly estimated, with the meta-analysed average shown in filled bars. The null expectation, equal to the percent of SNPs in each category, is shown by dashed, unfilled bars, with p-value reporting the difference from this expectation. Fold-enrichment relative to the null expectation shown in paranthesis below each category label. Left panel shows results from analyses of genotyped SNPs only, right panel shows analysis of genotyped and 1,000 Genomes imputed SNPs. Error bars define 95% confidence interval.

We evaluated the robustness of the variance-component estimates to deviations from model assumptions and differences between functional categories (see Methods, Supplementary Note). In particular, DHS variants harbor the smallest amount of LD and coding variants exhibit the greatest average evolutionary constraint (Table S2), motivating us to perform simulations using real genotypes and functional categories. The 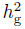 estimates were unbiased for simulated phenotypes with causal variants sampled uniformly (Fig. S2); entirely from a single category (Fig. S3); or with enrichment at DHS and coding mimicking the real data (Fig. S4). More extreme disease architectures with common, low-frequency, and category-specific causal variants yielded estimates that were nearly unbiased (Fig. S2, Fig. S5). We considered the possibility that causal variants lie just outside DHS, but our simulations argue against this possibility (see Methods, Fig. S6). An experiment in which the standard error was estimated via weighted jack-knife across chromosomes (which is conservative, reflecting genuine differences across chromosomes) yielded a slightly higher standard error and slightly lower DHS % 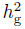 (71% s.e. 7.7%), but the enrichment was still highly significant (*P* = 5.5 × 10^−13^) and the overall results not substantially affected (Table S8, S9). The impact of shared controls on estimated standard errors, quantified by empirical resampling, was around λ_GC_ = 1.4 for most categories, yielding adjusted standard errors that remained significant for all categories except UTR (Table S10). Finally, accounting for case-control ascertainment using Haseman-Elston regression-based estimates^21–23^ of 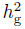 yielded no significant differences in 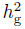 (Table S11, S12).

### Partitioning 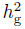 within DHS regions

We further partitioned DHS into functional sub-categories, assessing significance relative to all DHS. We used ENCODE-ChromHMM classifications of enhancer regions^24^ to partition DHS (15.7% of the genome) into those that overlap predicted enhancers (3.2% of the genome) versus other DHS (Fig. 2A). The enhancer-DHS explained 31.7% (s.e. 3.3%) of total 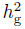, yielding an enrichment of 9.8× vs. all SNPs (1.9× vs. all DHS; *P* = 5.1 × 10^−4^). We partitioned DHS into regions that were called in two or fewer cell-types (“specific” after merging similar tissues) versus others (Fig. 2B). We observed a significant enrichment for cell-type specific DHS (6.1× vs. all SNPs; 1.3× vs. all DHS; *P* = 3.2 × 10^−3^). The enrichment was not significant when repeating this analysis for enhancer and non-enhancer DHS separately. We split the DHS into SNPs overlapping/non-overlapping the ENCODE database of DNaseI Digital Genomic Footprinting (DGF) regions (8.5% of the genome), which are expected to precisely map sites where regulatory factors bind to the genome^25^ (Fig. 2C). We observed no difference in 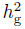 relative to other DHS sites (1.0×, *P* = 0.90). However, DGF annotations were collected for only a subset of DHS cell-types analyzed, and analysis in additional cell-types is needed. Lastly, we partitioned the 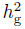 using an expanded DHS annotation (including regions overlapping coding, UTR, and promoters) into the remaining five major categories (Table S13), which yielded enrichment at DHS-coding of 34.4× vs. all SNPs (5.3× vs. all DHS, *P* = 1.35 × 10^−3^) and DHS-promoter variants of 13.2× vs. all SNPs (2.3× vs. all DHS, *P* = 7.90 × 10^−3^). Notably, unlike the non-DHS introns, we did not observe significant depletion at DHS-introns (0.9× vs. all DHS, *P* = 0.037). We observed nominally significant negative 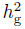 at the remaining non-DHS category (*P* = 0.002), likely due to underestimating standard errors in the analysis, and so treat the within-DHS enrichments as suggestive.

**Figure 2.**
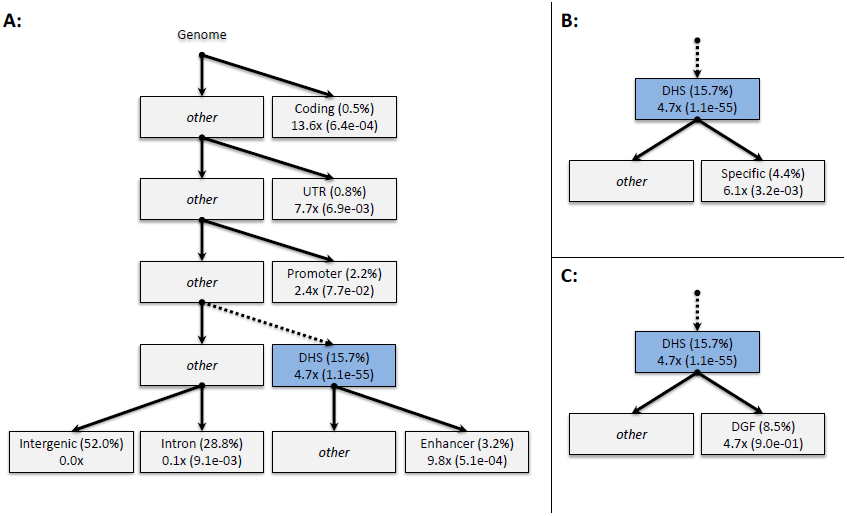
Hierarchical analysis of functional enrichment. DHS variants were further partitioned into three sub-categories: predicted enhancers, cell-type specific, and digital genomic footprinting (DGF) targets. Each block contains: on top line, the functional category and fraction of genome (in parenthesis); on bottom line, fraction enriched relative to the rest of the genome and p-value of enrichment relative to parent category (in parenthesis). DHS enrichment of 4.7× non-significantly different from 5.1× in Figure 1 due to additional free parameters.

To investigate the role of specific cell-types, we separately estimated enrichment in 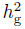 for DHS in each of 83 cell-type groups. For each trait and cell-type, we estimated 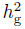 jointly from three components corresponding to DHS observed in that cell-type, other DHS not observed in that cell-type, and all other SNPs; assessing enrichment relative to all DHS. Based on our previous observation of heterogeneity, we performed meta-analyses across the six autoimmune traits (with MHC excluded) and across the five non-autoimmune traits. We observed six cell-types that were significantly enriched in autoimmune traits (conservatively adjusting for 83 tests, though the cell-types are highly correlated), and none significantly enriched in non-autoimmune traits (Table 1). Four of these six cell-types have previously been implicated in autoimmune diseases: Trynka et. al.^4^ identified GWAS hits for rheumatoid arthritis to be enriched within H3K36me3 peaks from CD8+ primary cells (at *P* = 0.0042); and Maurano et. al.^5^ identified nominally significant SNPs in a GWAS of Crohn’s disease to be enriched within DHS peaks from *T_H_* 1 T-cells, and nominally significant SNPs in a GWAS of multiple sclerosis to be enriched in DHS peaks from B-lymph cells. The remaining two significant cell-types, leukemia and fetal pelvis cells, are novel (additional nominally significant cell-types listed in Table S14).

**Table 1.** Cell-type and phenotype specific DHS enrichment. Fold-enrichment of 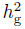 reported for cell-type-DHS observed as significant (after adjusting for 83 cell-types tested). Enrichment was measured in comparison to 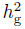 at DHS regions, to account for the background DHS enrichment. Results shown separately from meta-analysis of 6 autoimmune traits and 5 non-autoimmune traits. Instances where enrichment was also observed in Trynka et al. 2013^4^ or Maurano et al. 2012^5^ are indicated.

| Tissue type | Cell type | Autoimmune traits | Non-autoimmune traits | Published |
| --- | --- | --- | --- | --- |
| Blood | CD8<sup>+</sup> Primary Cell | 4.0 (1.7e-04) | 0.9 (7.9e-01) | <sup>4</sup> (RA) |
| Blood | Leukemia Cells | 3.5 (5.9e-05) | 0.9 (4.5e-01) |  |
| Blood | Lymphoblastoid Cell | 3.4 (3.1e-05) | 1.2 (8.9e-01) | <sup>5</sup> (MS) |
| Blood | Primary Th1 T-Cell | 5.8 (1.4e-05) | 1.8 (3.5e-01) | <sup>5</sup> (CD) |
| Bone Marrow | Monocyte CD14<sup>+</sup> | 4.3 (4.3e-04) | 1.1 (8.2e-01) | <sup>5</sup> (MS) |
| Fetal Kidney | Fetal Right Renal Pelvis | 5.4 (4.3e-04) | 0.9 (8.2e-01) |  |

### Heritability of functional categories in PGC and exome chip schizophrenia cohorts

We replicated our functional enrichment results in an independent cohort of 5,073 schizophrenia cases and 6,605 controls from the Psychiatric Genomics Consortium (PGC), imputed to 1,000 Genomes SNPs^26^. The WTCCC2 analysis restricted to schizophrenia produced a non-significant DHS enrichment of 2.6× (s.e. 1.47, *P* = 0.28) and intergenic 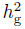 of 0.4× (s.e. 0.27, depletion *P* = 0.02) (Table S5). In the PGC data, the DHS enrichment was significant at 3.0× (s.e. 0.64, *P* = 1.9 × 10^−3^) and the intergenic category was significantly depleted at 0.4× (s.e. 0.12, depletion *P* = 3.6 × 10^−7^). The consistency of WTCCC2 and PGC enrichments indicates that platform artifacts are unlikely to be a major confounder.

We further analyzed an independent cohort of 2,500 schizophrenia cases and 3,875 controls of Swedish origin typed on both GWAS and exome chips (see Methods), to investigate the possible contribution of rare coding variants to missing heritability^27^, defined as the gap between our genome-wide estimates of 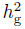 and the total narrow-sense heritability. The exome chip variants were primarily rare, consisting of 18% singletons and 64% non-singletons below 1% MAF. Due to the abundance of rare variants, we performed additional simulations to assess bias and demonstrated that frequency-stratified^28^ and LD-adjusted^16^ components provided unbiased estimates of 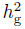 under a wide range of disease architectures (see Methods, Supplementary Note). We partitioned the heritability explained by GWAS chip and exome chip data into three separate variance components: non-coding, rare coding (MAF < 1%), and common coding variants. This partitioned analysis identified a total 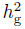 of 0.079 (s.e. 0.034) from all coding variants (Table 2), with only the 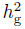 of 0.042 (s.e. 0.017) from common coding variants being significantly different from zero (*P* = 7.7 × 10 *^−^*^3^; rare coding *P* = 0.10). The 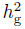 from rare variants remained non-significant even after partitioning according to PolyPhen score^29^, restricting to putative schizophrenia genes, or collapsing variants by gene to reduce the standard error (see Supplementary Note). Moreover, the estimate of DHS enrichment was unaffected by the inclusion of rare coding variants (Table S15), confirming that DHS enrichment was not an artifact of untagged coding variation in this cohort. We caution that our exome chip results pertain to rare variants included in the chip design (ascertained from 12,000 samples), but do not extend to extremely rare variants. However, our findings are consistent with a recent analysis of exome-sequencing data in schizophrenia, which identified a significant but modest (0.4%-0.6% of total variance) burden from rare coding variants in a subset of ~ 2500 genes^30^.

**Table 2.**
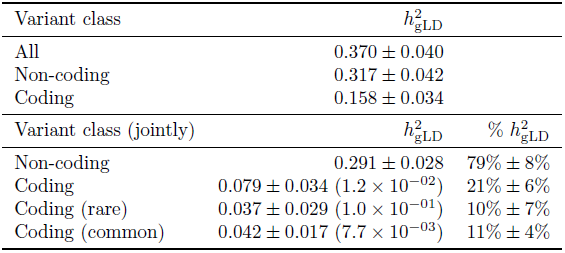
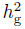 of schizophrenia from exome chip. Estimates of 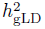 are reported from variance components (adjusted for LD) in the homogenous Swedish sub-population. Top panel shows estimates that include tagging of variants in other classes. Bottom panel shows joint estimates accounting for tagged variance due to LD. P-Values from a likelihood ratio test shown in parentheses.

### Limitations of current approaches to assess enrichment using summary statistics

Surprisingly, the DHS category was not substantially enriched in individual genome-wide significant SNPs identified in this data (0.91×, Table S16) nor in the full distribution of association statistics (Fig. S7). Several methods have been proposed for assessing functional enrichment based on GWAS summary statistics^5,9^, and we sought to evaluate their performance here. Using our observed genome-wide estimates of functional 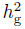, we simulated quantitative traits in a combined WTCCC2 cohort of 33,000 samples with 8,300 causal variants (corresponding to a large GWAS of a polygenic trait^31^), where 79% of heritability was explained by imputed DHS SNPs, 8% by imputed coding SNPs, and the remainder uniformly drawn from the other variant categories. Applying the variance-component strategy to the simulated cohorts correctly recovered the enrichment factors (Fig. 3A). We then conducted a standard GWAS on the simulated traits and plotted functional enrichment using two published strategies: stratified QQ-plots with variants classified based on aggregate LD with each category^9^; and p-value enrichment, comparing the fraction of significant SNPs in a category to the genome-wide average^5^ (see Supplementary Note). Both strategies yielded enrichment at coding variants through the full distribution of association statistics (Fig. 3B,C). However, proximal categories such as UTR and promoter, which were truly depleted, also appeared as enriched through tagging of significant coding variants. DHS variants were the least enriched non-intergenic category by both methods even though they made the single largest contribution to heritability. The p-value enrichment plot, in particular, demonstrates substantial enrichment at all other non-intergenic categories but little for DHS (Fig. 3B). This is likely due to lower power to detect DHS SNPs due to their lower average effect-size (relative to coding) and less LD. These simulations demonstrate that GWAS p-values, while partially informative, have a noisy relationship with the underlying category-specific heritability, motivating additional work on methods that can produce robust estimates of enrichment from summary statistics.

**Figure 3.**
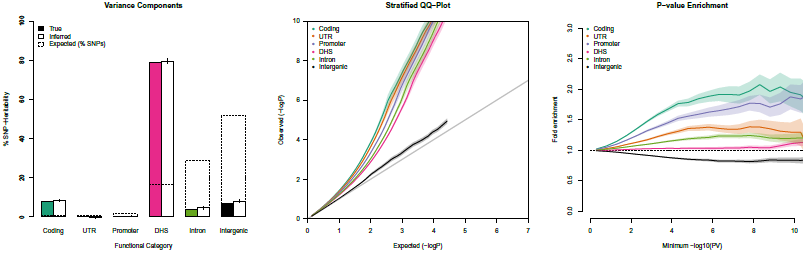
Enrichment from GWAS summary-statistics. We simulated traits where DHS and coding variants explain 79% and 8% of 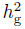, respectively, and computed GWAS statistics in a cohort of 32,000 samples. Enrichment in each functional category was measured using variance-components (left) and two previously described methods that rely on genome-wide association statistics (center, right). Though coding regions were identified as enriched, other categories exhibited false enrichment while DHS was incorrectly identified as least enriched by both methods. Shaded region and error bars represent standard error from 50 replicates.

To investigate whether the enrichment at significant loci was consistent with the genome-wide estimates, we partitioned 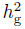 from SNPs lying within 1Mb of published GWAS loci for each trait (see Web Resources) (Fig. S8). The DHS enrichment of imputed SNPs at these known loci was highly significant (3.6×, *P* < 1 × 10^−20^) and was significantly higher than the DHS enrichment for genotyped SNPs only (*P* = 7.9 × 10^−18^ for difference). A nominally significant difference was observed between the imputed enrichment at known loci and genomewide (*P* = 0.003 after correcting for shared controls) suggesting that large-effect loci may be somewhat less enriched.

### Fine-mapping analysis with functional priors

Estimates of functional 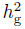 enrichment can guide fine-mapping analysis, where the goal is to identify a minimal set of SNPs that includes the underlying causal variant(s)^32,33^. Here, we applied the function-specific estimates of enrichment as priors for fine-mapping in four traits (RA, SP, T2D, CAD) with imputed summary statistics publicly available (see Web Resources). Given that all SNPs at genome-wide significant loci in these cohorts explain only 1.5% of the trait variance on average, we do not expect possible partial sample-overlap with WTCCC data to be a significant confounder. Though fine-mapping analysis ideally involves targeted sequencing or genotyping,^32^ observed that the latter had little impact on their fine-mapping analysis when compared to imputed data, so we expect imputed markers to be a reasonable proxy. For each trait, we restricted the analysis to loci that were genome-wide significant in these data (to ensure adequate signal for fine-mapping) and computed the 95% credible set^32^ of causal SNPs at each locus with and without applying functional priors (see Methods). On average, we found that the six main functional priors reduced the credible set of causal variants by 30% across the four traits (Table 3). The largest reduction of 63% was observed in RA, where the total credible set for 5 loci (MHC excluded) was reduced from 69 SNPs to 26. For comparison, including only coding variant enrichment as a prior reduced the credible sets by 5% on average and had no reduction for RA.

**Table 3.** Credible sets of causal SNPs at known associated loci. For each trait, genome-wide significant loci from meta-analysis association statistics were reduced to 95% credible sets with and without functional priors. Right-most four columns describe number of SNPs in credible set obtained from each prior type (with % relative to flat prior in parantheses). “Flat prior” corresponds to standard analysis with no functional information; “Coding prior” uses only enrichment at coding variants; “Main func. priors” includes all six priors from the main functional analysis; “Main & enh. priors” uses all six main priors and the enhancer-DHS prior.

| Phenotype | # Loci | Tot. SNPs | Flat prior | Coding prior | Main func. priors | Main & enh. priors |
| --- | --- | --- | --- | --- | --- | --- |
| RA | 5 | 8393 | 69 | 69 (100%) | 26 (37%) | 26 (37%) |
| SP | 11 | 84684 | 686 | 641 (93%) | 564 (82%) | 531 (77%) |
| T2D | 13 | 24799 | 101 | 90 (89%) | 84 (83%) | 83 (82%) |
| CAD | 16 | 27685 | 112 | 112 (100%) | 90 (80%) | 86 (77%) |
| CAD (metabo) | 34 | 7498 | 325 | 325 (100%) | 264 (81%) | 260 (80%) |

### Polygenic risk prediction

The 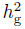 value for a set of SNPs is an upper-bound on the prediction accuracy of a polygenic score constructed from those SNPs^34–36^. To evaluate the impact of functional partitioning on risk prediction, we compared GBLUP^37^ prediction accuracy using six jointly estimated functional components vs. a single genome-wide component in the PGC schizophrenia data (see Methods). Of the six components, DHS yielded the highest individual *R*^2^ (0.055) and coding yielded the lowest (0.009) (Table S17). A single degree of freedom GBLUP prediction from the sum of all six components yielded a highly significant *R*^2^ of 0.061 (*P* < 10^−20^). However, GBLUP prediction using a single component was only slightly less accurate, with *R*^2^ = 0.058 (*P* = 2.6 × 10^−7^ for difference). Though highly statistically significant, this difference is substantially lower than would be expected in the case of independent markers, due to a high degree of correlation across components (see Supplementary Note).

## Discussion

In this study, we quantified the contribution to heritability of variants in diverse functional categories, implicating regulatory regions marked by DHS as explaining an average of 79% of 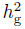 across 11 traits. We demonstrated by extensive simulations that our variance-component strategy yields robust estimates that account for LD between categories. In a schizophrenia cohort of 6,375 samples, the contribution from rare, exome-chip variants was non-significant and did not impact the observed DHS enrichment.

With genome-wide association studies identifying primarily noncoding variants, many hypotheses have been developed to explain the architecture of complex traits, including noncoding RNA, DNA methylation, alternative splicing, and un-annotated transcripts^10,38^. Several previous studies have demonstrated an excess of significant GWAS associations in regulatory categories. In particular,^39^ observed 2× enrichment in cell-type relevant enhancers;^40^ identified 1.12× enrichment at DHS; and^5^ identified 1.4× enrichment at DHS as well as enrichment in cell-type relevant DHS. However, we have inferred biological function from all SNPs simultaneously instead of one GWAS hit at a time. Our findings constrain most of 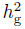 to the 16% of SNPs that lie in DHS, particularly in enhancers, and suggest that the other proposed mechanisms are unlikely to make substantial independent contributions. Furthermore, an analysis of DHS regions narrowed to cover 1% of the genome still explained 20% of 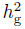 directly (and 61% in total), potentially motivating a DHS-targeted genotyping chip analogous to the exome chip^41^. More generally, our approach provides a means of assessing biological hypotheses of contributions to disease heritability.

Quantifying functional enrichment also has direct implications for GWAS and fine-mapping. Known functional priors can be used to optimize fine-mapping studies^32,42–45^ and the enrichments we observed substantially reduced the set of potential causal variants in four traits. On the other hand, despite the appeal of analyses of summary statistics in many contexts^46–49^, published methods for assessing enrichment from genome-wide summary statistics are less accurate than variance component methods, and new methods are needed for this type of data. This may explain why methods that use GWAS summary statistics from functional categories to inform association studies have yielded modest increases in power^33^. Surprisingly, the improvement in risk prediction from functionally stratified components was limited due to pervasive LD across categories.

A limitation of assessing enrichment from GWAS platforms is that it cannot account for untagged causal variation, representing roughly half of total narrow-sense heritability^50^. While we have shown that rare coding variants are unlikely to alter the DHS enrichment, the missing heritability may lie in other categories. The precision of inferred enrichment is also limited by the underlying annotations. We have shown that sub-annotation of DHS regions can yield additional enrichment, and it is likely that other functional categories — including additional epigenetic marks and cell-types — will be highly informative. Finally, multi-trait meta-analysis limits precision due to differences between traits, but larger sample sizes will enable conclusions specific to individual diseases.

## Online Methods

### Variance-components approach for estimating 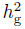

For a single component of genotyped SNPs we define 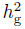, an underlying parameter in the population, as the *r*^2^ resulting from the best linear prediction of genotyped SNPs. Thus, 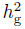 represents the amount of phenotypic variance explained by the genotyped SNPs and any linked SNPs. When SNPs are independent, this definition extends naturally to multiple components, with 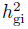 for component *i* defined as the 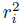 from SNPs used to construct that component. When SNPs are no longer independent (as in GWAS data) this definition remains valid as long as the individual SNP effect-sizes are i.i.d, which we expect to be approximately the case in polygenic traits with thousands of underlying causal variants.

Formally, for *a* functional categories each containing the set of SNPs defined by *M_i_*, we model the phenotype as a sum of individual SNP effect-sizes 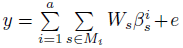 where the effect-sizes are drawn from individual normal distributions 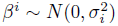 and 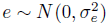. We assume that for each annotation *i*, SNPs normalized to have mean 0 and variance 1 are contained in the matrix *W_i_*. The variance of the phenotype is then modeled as 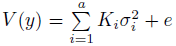 where each *K_i_* represents a genetic relationship matrix computed directly from the SNPs in annotation *i* as 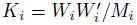. The corresponding *σ* are then jointly inferred using the REML algorithm in GCTA^11,14^, yielding 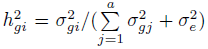. The inverse of the final Average Information matrix yields an estimate of the corresponding error covariance matrix of the variance-component estimates^51^. We use the error-covariance matrix and delta method^52^ to compute standard-errors on 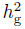 and % 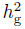 while accounting for error correlations (referred to here as the “analytical” standard error). All estimates were transformed to the liability-scale using prevalence values in Table S3.

### Functional categories

DNase-seq libraries from ENCODE and Epigenome Roadmap projects were downloaded in May 2012. Biological replicates were merged into a single library (GEO accession numbers available in Supplementary Table 30). Raw read sequences were aligned to hg19 using BOWTIE^53^ v1.0. DHS peaks were called using MACS v2.0 with FDR <0.01. For the primary analysis, all peaks were merged into a single DHS annotation. The resulting annotations are available for download (see Web Resources). The 98% of the primary annotation was covered by the DHS regions analyzed in Maurano et. al.^5^, and 67% was covered by the DHS regions analyzed in^3^ by 67%. For the cell-type analysis, duplicate lines were merged to form a final set of 83 unique cell-types.

Combined chromHMM genome segmentations were downloaded for 6 cell-lines (see Web Resources). All regions classified as WE (weak enhancer) or E (enhancer) were then combined into a single enhancer annotation.

DNaseI Digital Genomic Footprinting (DGF) regions were downloaded for 57 cell-lines (see Web Resources). All regions from the narrow-peak classification were then merged into a single DGF annotation.

### Simulations of partitioning 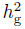 by functional categories

As has been shown previously, a relationship between minor allele frequency and effect-size can bias the 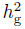 estimate by over-representing certain SNPs^16,28,54^. We performed simulations in genotyped and imputed data to confirm that estimates of enrichment are well-calibrated. We considered three disease-architectures: MAF-independent, where SNPs with any MAF can be causal; low-frequency, where only SNPs below 0.05 MAF can be causal; and DHS-low-frequency, where causal DHS variants are drawn from MAF below 0.05 and all other variants are drawn from any MAF. The low-frequency and DHS-low-frequency architecture represent the greatest possible extremes of MAF-dependence or functional category-specific MAF-dependence.

For each disease-architecture and simulation, causal variants were randomly sampled such that 7% of the SNPs were causal (~10,000 SNPs in genotyped data) and normally distributed effect-sizes were applied to construct additive quantitative traits with total 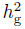 of 0.5 (such that each SNP explains equal variance in expectation). By sampling equal fractions of causal variants from each functional category, we expect to see no significant 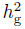 enrichment relative to category size. We performed the simulations using real genotypes from the WTCCC1 CAD samples (having roughly average sample size), with 1,000 randomized trials for each disease-architecture. As expected, no enrichment was observed under the MAF-independent architecture for either genotyped or imputed markers (Fig. S2). Under the low-frequency and DHS-low-frequency disease architectures, slight but statistically significant deviations were observed, primarily for the promoter and UTR groups which are very small and in tight LD with each other (Fig. S2). Next, we considered two contrived annotations constructed from either the 16% of SNPs with most LD partners and the 16% of SNPs with fewest LD partners, to mimic a high/low LD category approximately equal in size to DHS. Testing the uniformly drawn MAF-independent architecture, we again observed no enrichment for either the high-LD (1.02× se 0.01) or low-LD (1.02× se 0.03) annotations over 1,000 trials. Lastly, we considered a realistic MAF-independent disease architecture where causal variants were drawn from imputed DHS and coding categories with 79% and 8% (respectively) and all other categories uniformly mimicking the enrichment we observed in real data. We simulated 100 phenotypes from this architecture in a 33,000 sample union of the imputed WTCCC2 individuals, and the variance-components accurately recovered the corresponding enrichment (Fig. 3A). In this realistic scenario, power to detect significant DHS enrichment (*P* < 0.05) surpassed 95% at a sample size of 7,000 (Fig. S9).

Focusing on the imputed SNPs, we considered the other end of the enrichment spectrum, where all causal variants are drawn from a single functional category. In total, we evaluated 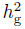 accuracy in 200 × 6 simulations for each of the MAF-independent and low-frequency architectures. In the MAF-independent simulations, we again observed unbiased estimates of 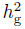 around 1.0 for the true causal category and 0.0 for all others (Fig. S3). In the low-frequency causal simulations, we observed nearly unbiased estimates of 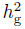, with slight but significant inflation at coding (1.2×) and UTR (1.07×) categories (Fig. S5). Lastly, we tested a deliberately severe architecture where causal variants are sampled from intronic and intergenic regions directly adjacent to DHS (0-500bp or 500bp-1000bp of a DHS boundary). In this scenario, we may expect some non-zero DHS 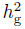 due to causal non-DHS SNPs in perfect LD with DHS SNPs. The DHS component picked up 50% (0-500bp) and 20% (500bp-1000bp) of the non-DHS 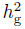 (Fig. S6), indicating that DHS 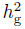 may still include some tagged variance from SNPs very close to the DHS boundary. However, the fact that 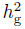 of intronic and intergenic regions is not significantly different from zero in real data implies that this effect is expected to be minor.

To investigate the differences between genotype and imputation-based estimates in real data, we partitioned 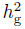 in the low-frequency, category-specific imputed phenotypes using components from genotyped SNPs only. If the genotypes are reasonable proxies for imputed variants, 100% of 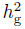 should again be partitioned into each focal category. Instead, we observed significant deviations for all of the categories, with 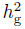 falsely partitioned into nearby categories due to incomplete tagging (Fig. S10). In particular, less than half of the 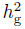 at DHS is correctly partitioned into the DHS category, consistent with the differences between 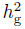 partitioned from genotyped and imputed SNPs in real data.

### WTCCC and WTCCC2 data sets spanning 11 diseases

We analyzed 7 traits from the WTCCC1 and 4 from the WTCCC2 for a total 47,000 samples. Estimates of 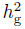 are particularly sensitive to individually small artifacts/batch-effects^13,55^. We followed the rigorous QC protocol outlined previously^16^: removing any SNPs that were below 0.01 minor allele frequency, above 0.002 missingness, or had deviation from Hardy-Weinberg equilibrium at a p-value below 0.01. For each case-control cohort we removed SNPs that had differential missingness with p-value below 0.05. We excluded one of any pair of samples with kinship entries ≥ 0.05^13^ and performed five rounds of outlier removal whereby all individuals more than 6 standard deviations away from the mean along any of the top 20 eigenvectors were removed and all eigenvectors recomputed^56^. For all autoimmune diseases analyzed (RA, CD, T1D, UC, MS, AS) we also exclude from the analysis any SNPs in the region around the MHC locus (chr6:26-34Mbp), which has been repeatedly documented to have a complex LD structure and many heterogeneous variants of strong effect for these traits.

All reported meta-analysis values were computed using inverse-variance weighting to account for different levels of error across 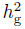 estimates. This estimate is expected to be tilted towards the autoimmune traits, which made up 6/11 traits analyzed as well as 20,461/30,158 cases analyzed.

### Imputation

The WTCCC1 samples were phased and imputed as described in^16^. The WTCCC2 samples were split into two cohorts by platform and each cohort imputed separately with the following protocol. All samples in a cohort were phased together in 10Mbp blocks using HAPI-UR^57^ (see Web Resources), with three rounds of phasing and consensus voting. All phased samples in a cohort were then imputed in 1Mbp blocks using IMPUTE2^58^ (see Web Resources) and the 1000 Genomes^59^ Phase I integrated haplotypes (9/2013 release; see Web Resources) with no singletons. Where relevant, the Oxford recombination map^60^ was used. Markers with an INFO score greater than 0.5 were retained. Finally, SNPs were excluded if they met any of the following criteria in any case or control population: Hardy-Weinberg p-value < 0.05; per-locus missingness > 0.05; minor allele frequency < 0.01; case-control differential missingness p-value < 0.05.

### PGC and exome chip schizophrenia cohorts

For estimates of rare variant 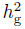, we analyzed 12,674 samples of Swedish origin typed on GWAS and exome chips. The exome chip yielded 238,652 SNPs (including monomorphic sites) of which 10,567 were also typed on a mix of Affymetrix GWAS chips (exome chip calls retained). The GWAS chip data contained an intersection of 163,051 SNPs typed on all platforms as well as per-platform imputation from the 1,000 Genomes with a total of 5,053,934 SNPs imputed on all platforms. Principal component analysis of the GWAS data revealed a large cluster of “homogenous” Swedish samples as well as clines related to Northern Swedish and Finnish admixture (Fig. S11). After excluding samples that were not typed on both GWAS and exome chip, failed QC, were PCA outliers by 6 standard deviations, or were in a pair with GRM values > 0.05 (close relatives) we retained a total of 8,967 samples of which 6,375 were of “homogenous” Swedish ancestry. The allele frequency classes of the exome chip data for these samples are outlined in Table S18. In all of our analyses, “rare variants” refer to MAF < 0.01 and “common variants” refer to MAF ≥ 0.01. All simulations were performed on the homogenous samples (without principal components) and all real data analyses additionally included the top 20 principal components as covariates (to account for any residual population structure).

### Unbiased estimates of 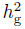 with rare and common variants

Unlike partitioned common variants, the 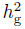 estimates from exome chip data could be substantially biased due to the abundance of rare variants^16,28,54^ (Fig. S12). We performed simulations across the full causal allele frequency spectrum and found that joint estimates from two frequency-stratified^28^ components computed from rare (MAF*≤* 0.01) and common (MAF> 0.01) SNPs eliminated most of the observed bias. Subsequently adjusting each component for LD (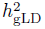, as described in^16^) completely eliminated bias for normalized effect-sizes, and yielded the most accurate estimate for standard effect-sizes (Supplementary Note, Fig. S13). We report estimates from joint components with LD adjustment (Table 2) and without (Table S19).

### Fine-mapping analysis with functional priors

For each genome-wide significant locus, we performed fine-mapping analyses using the previously described Bayesian model^32,33^. Each locus was defined as the union of 1MB windows around any SNP with p-value< 5 × 10^−8^. Summary statistics consisting of individual SNP effect-sizes and effect-size error were converted to Bayes factors as described in^33,61^ and multiplied by either a flat prior or the genome-wide functional prior (computed as the estimated 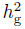 per SNP of the SNP category in the corresponding trait). The credible set for each locus was then computed for each scenario by including SNP Bayes factors from highest to lowest until the sum of Bayes factors in the set was at least 95% of the sum of the Bayes factors at the locus.

### Polygenic risk prediction

We analyzed 5,072 schizophrenia cases and 6,605 from the Psychiatric Genomics Consortium^26^. Imputation and quality control for these samples was described in^31^, yielding a total of 4,463,775 SNPs observed in all cohorts. BLUP coefficients were computed in GCTA^14^ (see Web Resources) using the imputed data in a model with a single genome-wide component and a separate model with the six functional category components and converted into SNP effects. Risk scores were then computed from the SNPs and effects for each corresponding component. We assessed prediction accuracy using 10-fold cross-validation, where component-specific heritability and BLUP values were only estimated in the ∽ 10,000 training samples. To account for population structure we included 10 principal components as fixed-effects in training the BLUP. We also included the same number of PCs when evaluating the predicted phenotype in a logistic regression with the true phenotype, reporting the Nagelkerke pseudo-*R*^2^ of each model minus that of the principal components.

## Acknowledgements

We thank Manolis Kellis, Abhishek Sarkar, Joe Pickrell, X Shirley Liu, Nick Patterson, Sara Lindstrom, Peter Kraft, and Shamil Sunyaev for helpful discussions, and Amy Williams for assistance with HAPI-UR. This research was funded by NIH grants R01 MH101244, R21 ES020754, R03 HG006731, NIH 1U01HG0070033, Doris Duke Clinical Scientist Development Award, and NIH fellowship F32 GM106584. We also acknowledge grant funding from ARC (DE130100614 and FT0991360) and NHMRC (613602 and 1050218)

**PGC SCZ Working Group**: S.R., B.M.N., S.M.P., B.J. Mowry, I. Agartz, F. Amin, O.A. Andreassen, M.H. Azevedo, N. Bass, D.W. Black, D.H.R. Blackwood, R. Bruggeman, N.G. Buccola, W.F. Byerley, W. Cahn, R.M. Cantor, K. Choudhury, S. Cichon, C.R. Cloninger, P. Cormican, A. Corvin, D. Curtis, S. Datta, S. Djurovic, G.J. Donohoe, J. Duan, F. Dudbridge, A. Fanous, R. Freedman, N.B. Freimer, M. Friedl, P.V. Gejman, L. Georgieva, I. Giegling, M. Gill, H. Gurling, L. De Haan, M.L. Hamshere, T.F. Hansen, A.M. Hartmann, P.A. Holmans, C.M.H., A. Ingason, A.K.K., R.S. Kahn, M.C. Keller, E. Kenny, Y. Kim, G.K. Kirov, B. Konte, L. Krabbendam, R. Krasucki, J. Lawrence, P.H. Lee, T. Lencz, D.F. Levinson, J.A. Lieberman, D.Y. Lin, D.H. Linszen, P.K.E. Magnusson, W. Maier, A.K. Malhotra, M. Mattheisen, M. Mattingsdal, S. Mane, S.A.M., A. McIntosh, A. McQuillin, H. Medeiros, I. Melle, V. Milanova, D.W. Morris, V. Moskvina, I. Myin-Germeys, M.M. Nthen, C. ODushlaine, A. Olincy, L. Olsen, R.A. Ophoff, M. J. Owen, C.N. Pato, M.T. Pato, B.S. Pickard, J. Pimm, D. Posthuma, V. Puri, D.J. Quested, H.B. Rasmussen, M. Rietschel, L. Rossin, D. Ruderfer, D. Rujescu, A.R. Sanders, T.G. Schulze, J. Shi, J.M. Silverman, D. St. Clair, T.S. Stroup, S. Thirumalai, J. Van Os, P.M. Visscher, T. Wassink, S. Zammit, P. Sklar, M.J.D., M.C. ODonovan, N. Craddock, P.F.S., K.S. Kendler

## Web Resources

1000 Genomes phase 1 reference panels: https://mathgen.stats.ox.ac.uk/impute/impute_v2.html#reference

CARDIoGRAM CAD summary statistics: http://www.cardiogramplusc4d.org/downloads/

DIAGRAM T2D summary statistics: http://diagram-consortium.org/downloads.html

DNaseI Digital Genomic Footprinting (DGF) annotations: http://hgdownload.cse.ucsc.edu/goldenPath/hg19/encodeDCC/wgEncodeUwDgf/

ENCODE-ChromHMM enhancer annotations: ftp://ftp.ebi.ac.uk/pub/databases/ensembl/encode/integration_data_jan2011/byDataType/segmentations/jan2

Exome chip design: http://genome.sph.umich.edu/wiki/Exome_Chip_Design

Functional annotations can be downloaded from: http://www.hsph.harvard.edu/alkes-price/software/

GCTA: http://www.complextraitgenomics.com/software/gcta/

GWAS catalog, downloaded on 2013.10.18: http://www.genome.gov/gwastudies/

HAPI-UR: http://genetics.med.harvard.edu/reich/Reich_Lab/Software.html

IMPUTE2: https://mathgen.stats.ox.ac.uk/impute/impute_v2.html

Oxford recombination map: http://hapmap.ncbi.nlm.nih.gov/downloads/recombination/

Rheumatoid arthritis summary statistics: http://www.broadinstitute.org/ftp/pub/rheumatoid_arthritis/Stahl_etal_2010NG/

Sweden+PGC1 schizophrenia summary statistics: https://pgc.unc.edu/Sharing.php#SharingOpp

